# Global slowdown in the rate of invasive alien species establishment

**DOI:** 10.64898/2026.08.21.746256

**Authors:** Melodie A. McGeoch, Rachel T. Mason, Saxbee Affleck, Jonathan Belmaker, Jean Cossi Ganglo, Walter Jetz, Rachel I. Leihy, Benjamin R. Shipley, Bharat Babu Shrestha, Wojciech Solarz, Marten Winter

## Abstract

The number of species introduced outside of their historical ranges by human activity continues to rise^1^. A subset of these species establishes, form self-sustaining populations, and some – invasive alien species – go on to cause substantial harm to biodiversity and ecosystems^2^. Preventing new invasive alien species from establishing is the key focus of interventions, because post-establishment management is costly and often fails^3,4^. However, it remains unclear how effective multilateral efforts have been in curbing the rise. Here we show that the emergence of new invasive alien species across countries is slowing, and trends are similarly negative across geographically diverse countries. Using data and modelling advances, we find a 35% reduction in the establishment of new invasive alien species over a policy-relevant 50-year time frame. The findings directly inform the assessment of progress for the invasive alien species target of the Kunming-Montreal Global Biodiversity Framework^5^, and provide a global baseline for monitoring rates of invasive alien species establishment. Furthermore, the slowdown suggests that policy and investment over recent decades to prevent invasive alien species from entering and establishing in countries have had a positive effect.

---

Invasive alien species (IAS) are a major direct driver of biodiversity loss, causing negative impacts on nature across all regions and taxa and contributing to 60% of recorded population extinctions^6^. The economic and social costs of biological invasions are substantial – conservatively estimated at tens of billions of US $ annually – and expected to multiply each decade^4,7^. The invasion process itself, driven by human activities and resulting in the transfer of species beyond their natural ranges, is now scientifically well understood^3,8,9^. The problem has also received growing policy attention for at least four decades^6,10^, with pervasive increases in the rate of biological invasion assumed^2^. Yet, it remains unclear whether the rate of IAS establishment is continuing to rise or perhaps beginning to slow in response to co-ordinated interventions internationally and nationally to prevent their introduction, establishment and spread. Answering this question is essential at a time when interactions among climate change, pollution, land-use change, and other drivers are amplifying invasion risks in ways that are difficult to predict and increasingly costly to manage^11^.

Global analyses document the accumulation of alien species since 1700 CE, with numbers of new records rising across taxa and regions from the mid-19th century through to the present^1,12^ with two distinct invasion waves (pre 1914 and 1960-present)^13^. An increase in the rate of biological invasion is the current baseline assumption of invasion policy^2,13^. Widespread policy attention to the impacts, drivers and management of biological invasions, and the first biosecurity protocols, emerged in the 1990’s following the development of invasion biology as a discipline in the 1980s^14^. Evaluating the effectiveness of this invasion policy and its regulations therefore requires a timeframe that is shorter than the long history of invasion records, and more targeted to the period over which contemporary policy has been implemented.

Research on global invasion trends has previously focused on all alien species, rather than on the far smaller, priority subset of IAS that are established and have negative impacts on biodiversity and that therefore warrant the most urgent management attention (Extended Data Table 1). This distinction matters for policy because prioritising management efforts for those species known to have negative impacts is most effective and efficient for mitigating the impacts of biological invasions^15^. Recognising this, Target 6 of the Kunming-Montreal Global Biodiversity Framework (KM-GBF) calls specifically for at least a 50% reduction in rates of IAS introduction and establishment by 2030 (https://www.cbd.int/gbf/targets/6). Two policy cycles ago, the IAS target of the Convention on Biological Diversity (CBD) was not met, i.e. to have *pathways of introduction of alien invasive species controlled, and management plans in place for major invasive alien species* (2010 Biodiversity Target Framework)^16^. The number of documented IAS was considered to be a significant underestimate, and at the time only half of the countries had IAS-relevant national legislation^10^. In the following cycle, Aichi Target 9 of the Strategic Plan for Biodiversity 2011–2020, by contrast, was judged to be partially achieved with progress made on identifying and prioritising IAS, but with no evidence of a slowing in the number of new introductions of alien species^17,18^.

In addition to a need for estimates of establishment trends for a priority species subset and over a policy-relevant time period, there is a need for trend estimates that consider the bias resulting from lags in the IAS detection process^5,13^. The phenomenon of a lag between the date of alien species establishment and the date of its first record (Extended Data Table 1) is widely recognised, with the extent of the lag influenced by both observation effort and the likelihood of IAS detection that increases after establishment as the species population grows and spreads^19^. The effect of this lag is that raw trends in numbers of species tend to be biased upwards in recent years, complicating the interpretation of establishment trends and the estimation of invasion rates^5,20,21^. This lag can bias establishment trend estimates and increase uncertainty when assessing policy effectiveness^5,22^.

To estimate global and country-level IAS establishment trends over the policy-relevant period 1970–2020, while appropriately accounting for observation effort, here we combined three primary data sources: (i) IAS checklists^23^ which identify IAS that are established and with evidence of negative impact in each of 191 countries, providing information on the overall detected magnitude of species with impactful invasions (Extended Data Table 1); (ii) years of first establishment for each IAS in each country^1,24^ adding a critical temporal dimension; and (iii) annual counts of new biodiversity occurrence records per country, providing an important proxy for observation effort (detail in Extended Data Fig. 1). These data were integrated and harmonised through an automated workflow with targeted manual quality control steps (Extended Data Fig. 1; Methods). To disentangle the underlying establishment rate from the confounding effect of imperfect and temporally variable species detection, we applied four trend estimation models that make progressively more realistic assumptions about the IAS detection process: (i) a Naïve model (assuming perfect detection), (ii) the Solow and Costello model and (iii) its Constant Detection variant (parametrising the detection lag), and (iv) a Sampling model that incorporates observation effort proxy directly (Extended Data Fig. 1; Methods)^25^. For country-level analyses, modelling was applied to the 85 countries (43% of all countries) with sufficient data completeness; the global time series incorporated data from 98% of all countries, including five case-study countries where the data were reviewed and augmented by national experts. Model selection and comparison across the four models enable estimation of the rate of change in IAS establishment in the presence of varying observation effort and imperfect detection (Methods).

This approach allows us to provide a first global estimate of IAS establishment rates focused on the priority subset of species with documented negative biodiversity impacts and accounting for bias resulting from the IAS detection process. We find that over the period 1970–2020 the global rate has declined, countering the prevailing view that IAS establishment rates continue to increase unabated^6^.

## Global decline in IAS establishments

We find a total of 5087 unique species that have been classified and listed as IAS in at least one country, which is 16% of the estimated number of established aliens globally (using a comparable number of globally established alien species^26^; Table 1). Of the global total, an expected 598 (∼12%) IAS emerged over the 1970–2020 period (‘the period’ from here), with an estimated average of 9.12 new IAS established per year (Table 1). Although there are more than twice the number of plant (*n* = 3546) than animal (*n* = 1541) IAS globally, more new invasive alien animals (63% of the total) emerged than plants for the period; with an estimated average of 5.73 animal species and 3.39 plant species per year (Table 1). With the overall number of native animal species worldwide at least 18 times higher than the number of native plants^27,28^, the potential for new invasive animal establishments may continue to exceed that for plants.

**Table 1.**
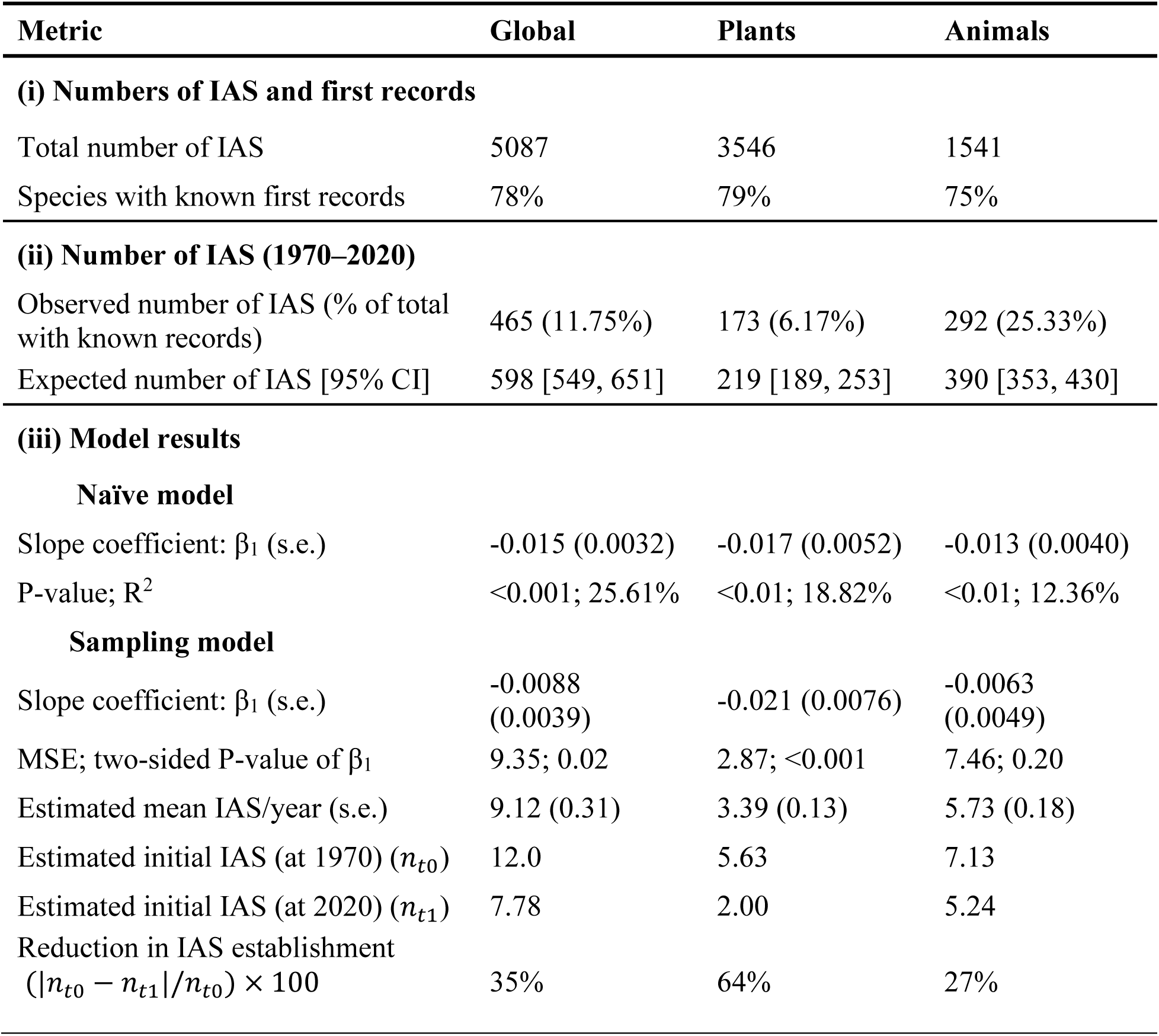
Global numbers and first records of established invasive alien species (IAS), overall (i) and for the period 1970–2020 (ii) . The expected number (ii) assumes a random temporal distribution of IAS before and after 1970 (Methods). (iii) Exponential model estimates of the rate of new IAS establishments (Methods). Results for both the Naïve model and the Sampling model (the best model in each case), and the % change in IAS establishment calculated from the sampling model estimates. MSE – mean square error.

At the global scale, the rate in the establishment of IAS has slowed over the period (β_1_ = -0.009, s.e. = 0.004) (Fig. 1A, Table 1 (iii, global slope coefficient)). The Sampling model was selected as the best model, but estimates of rates of change (β_1_) were consistent across the four models (negative in all cases with β_1_ ranging from -0.009 to -0.02) (Methods). This consistency across models with differing assumptions about the detection process strengthens confidence that the observed decline reflects a genuine reduction in establishment rate. The rate of the decline translates to a 0.88% decline in the number of IAS established globally per year. The estimated percentage reduction in IAS establishments over the five decades stands at 35%, with 12.0 species/year at the start and 7.78 species/year at the end of the period (Table 1, Sampling model). Rates for invasive alien plant and animal establishments were also negative (Fig. 1B- C; Table 1), consistently so across all models (Methods) with a stronger decline for plants (64% reduction) than for animals (27%; Table 1).

**Fig. 1.**
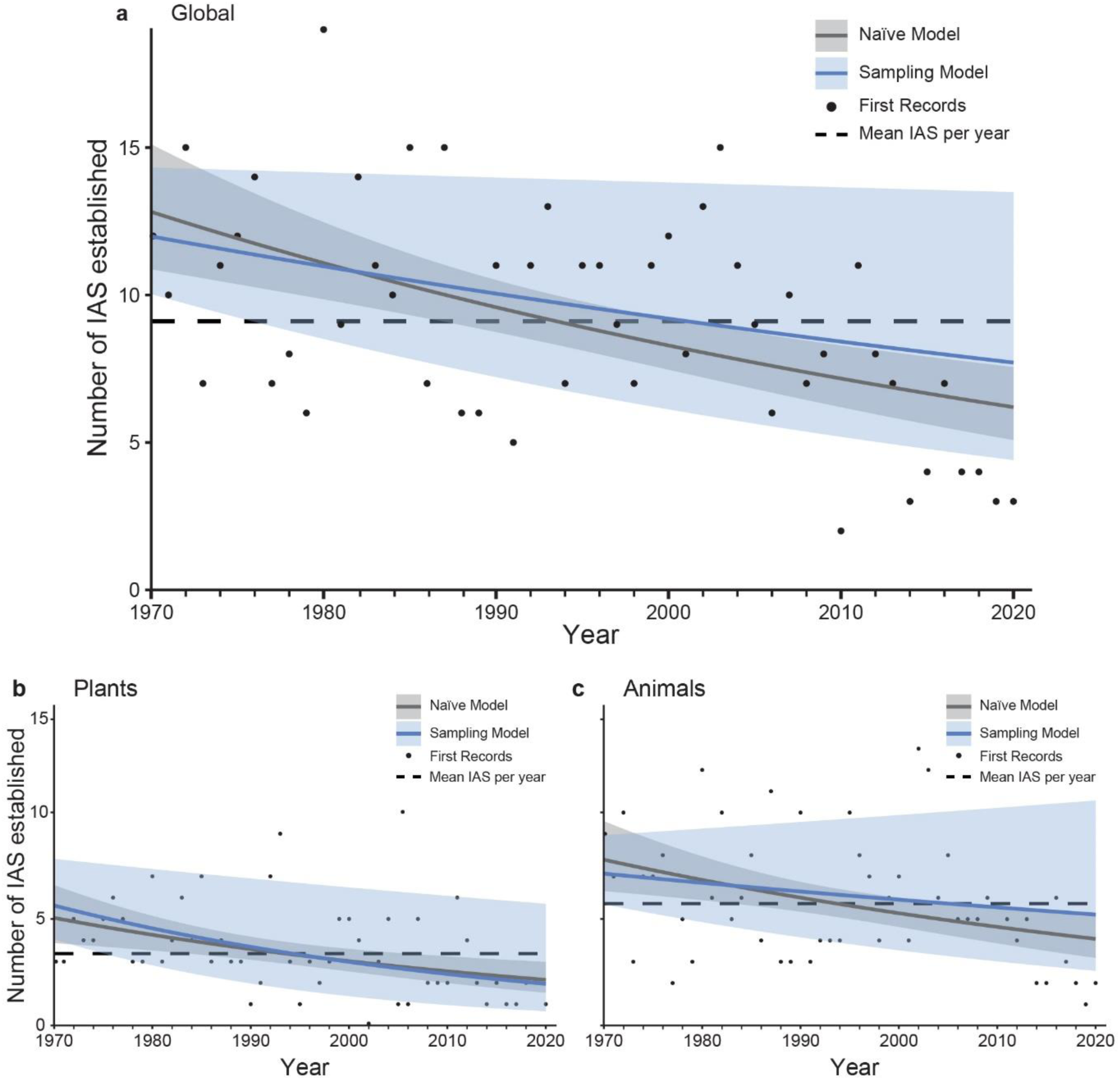
Global trend in the establishment of invasive alien species (IAS) across countries between 1970-2020. Total number (**a**) and for plants (**b**) and animals (**c**). The simplest (Naïve) model (grey) and the selected best model (Sampling model; blue) are shown in each case (Model results in **Table 1**). The ‘first record’ is the year in which the establishment of the species is recorded in any country for the first time. Mean IAS per year is the average number of IAS across the period 1970-2020. For consistency with the Naïve model, the predicted values of the establishment rate µ (defined by the β_0_ and β_1_ parameters) are plotted for the Sampling model (Methods). Error bars depict the ± 95 % confidence intervals around the mean predicted establishment rates for each of these exponential models, and the dashed black lines show the mean IAS establishments across the modelled period.

The observed slowdown is unlikely to be a consequence of declining observation effort ^13^. Using the external data proxy for IAS observation effort shows a strong growth in new biodiversity records over the period (P < 0.01, R² = 0.98; Methods). We make the assumption of a positive relationship between the trend in new biodiversity records per year for a country and the likelihood of new IAS being observed and recorded (Extended Data Fig. 1), While this is a proxy for IAS observation effort, it nonetheless suggests that the likelihood of newly established IAS being recorded has increased rather than decreased over the period. Furthermore, all four trend models, together designed to disentangle the establishment rate from the detection process, reproduce a negative trend that is robust to the timing and bounding of the policy-relevant period (Methods, Extended Data Fig. 2). The observed slowing in global IAS establishment rates therefore points to a mechanistic basis for the outcome rather than the effect of the detection process.

### Heterogeneity in country-level establishment trends

To examine whether the global slowdown is reflected at national scales, we estimated establishment trends individually for the 85 countries that met a first record data completeness threshold (Methods; Extended Data Fig. 1). More than three quarters of the countries modelled (78%) had either negative or flat establishment trends (Fig. 2A); 25% showed a negative trend in IAS establishment; trends were flat in 53% and increasing in 23% of countries (Fig. 2A). This remained the case even when only considering countries with flat trends where the average number of IAS establishments per year was less than the country-wide mean (64%; Fig. 2B), i.e. countries with a low and flat rate of IAS establishment are in a better position relative to those with flat but high rates of establishment. Overall, therefore, between 1970 and 2020 more of the countries examined had negative or flat trends in IAS establishment than positive trends.

**Fig. 2.**
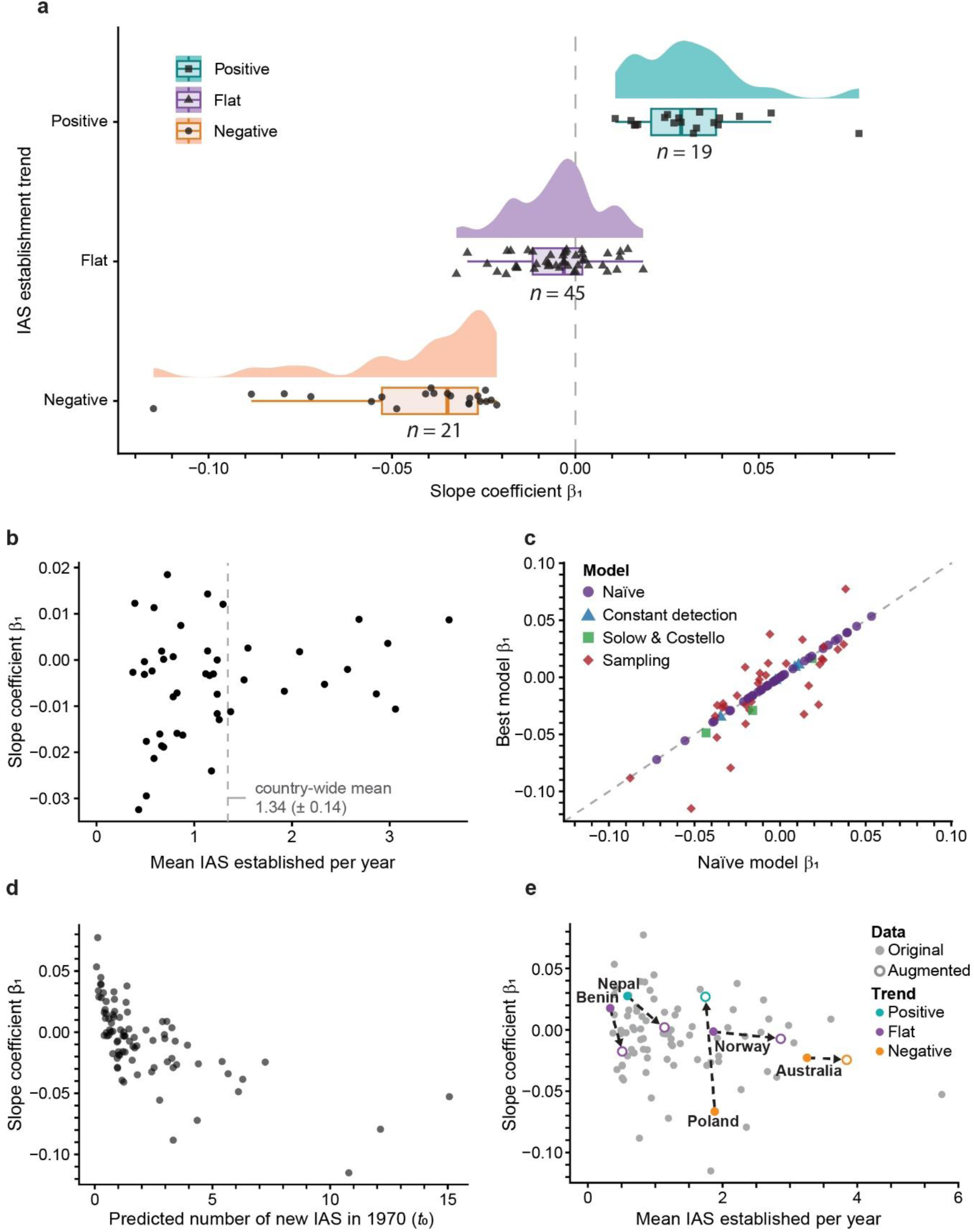
Variation in country estimates of invasive alien species establishment (1970-2020) (*n* = 85 modelled countries). (**a**) Per-country rates of invasive alien species (IAS) establishment (slope coefficient, β_1_ of the best model) for those with positive (+), flat and negative (-) trends. (**b**) For countries with a flat trend (*n* = 45), the relationship between the mean number of IAS established per year (best model per country) and the slope coefficient (β_1_) from that model relative to the country-wide mean. (**c**) Relationship between the slope coefficients (β_1_) of the Naïve model and the best model for each country (note that in some cases the Naïve model is the best model, Methods). (**d**) The relationship between the predicted number of new IAS in 1970 (*t*_0_) and the best model slope coefficient for all modelled countries. (**e**) The mean number of IAS established per year against the slope coefficient (β_1_) of the best model for each country (grey points, *n* = 85), highlighting changes in the trend and best model slope coefficients for five case-study countries where the data were augmented by country experts.

Country-level rates of change in the establishment of IAS varied widely geographically (Fig. 3; Extended Data Fig. 3D-F), from a high of an 8% increase per year in establishments (β_1_ = 0.077), to a low of a 12% decline (β_1_ = -0.115) in establishments per year (Fig. 2C; Extended Data Fig. 3). Across countries, the mode for those with negative trends was a 2.7% decrease in establishments per year, and a 2.9% increase per year for those with a positive trend (Fig. 2A). Countries with higher rates of establishment early in the period were more likely to show declining trends, whereas those with low early rates were variable but more likely to have positive IAS establishment trends (Fig. 2D).

**Fig. 3.**
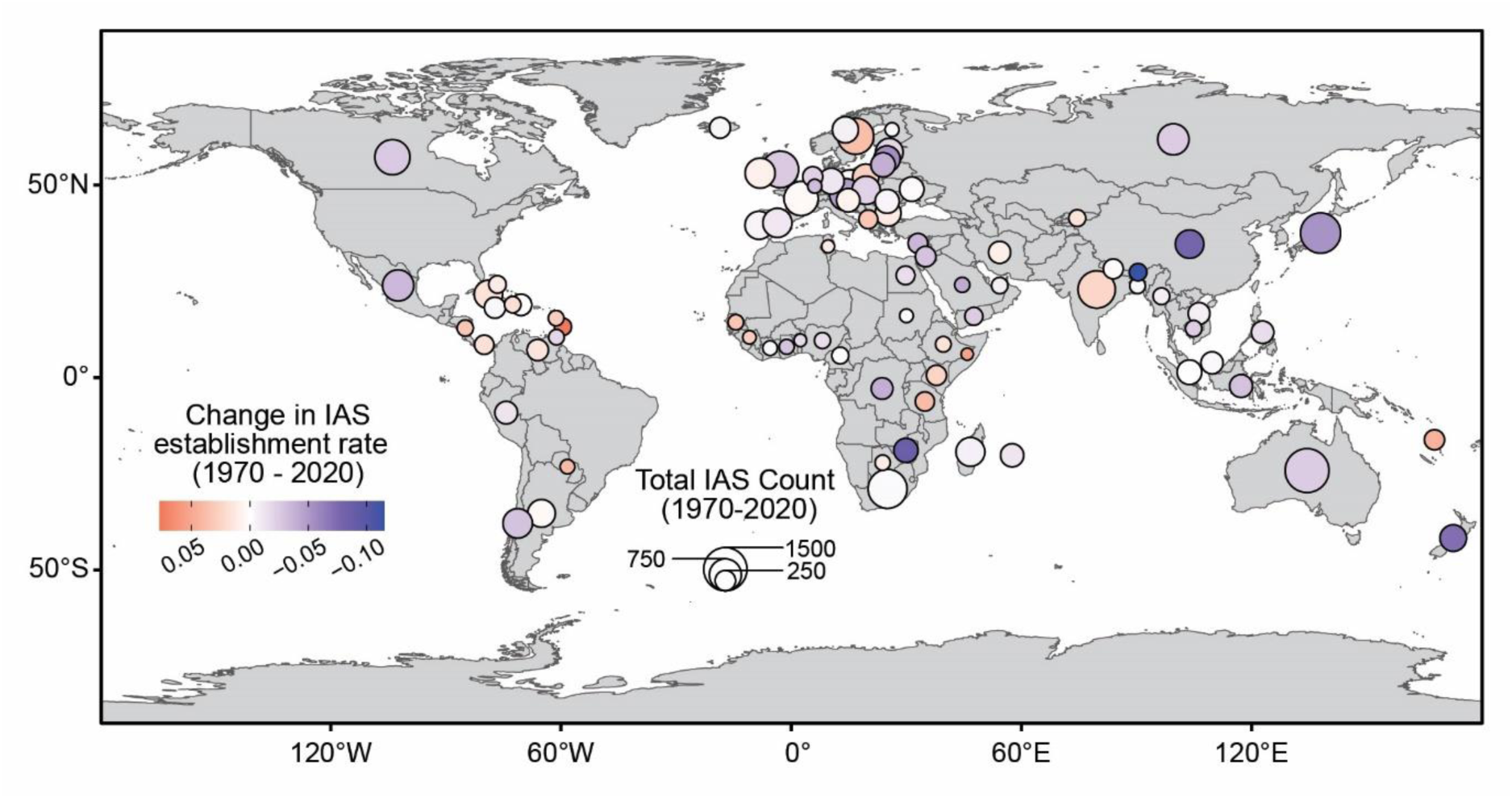
Patterns of invasive alien species (IAS) establishment, showing geographic heterogeneity in the distribution of total number of IAS (relative circle size) and rate of change (slope coefficient β_1_; circle fill colour) in IAS establishment for modelled countries (*n* = 85). Shaded countries encompass the data included in the global analysis (grey); circles are positioned over centroids of countries that were modelled individually.

The flat establishment trend outcome found in 53% of the cases (Fig. 2A-B) has two alternative implications for countries, depending on the average number of IAS established per year^5^. A flat trend with high annual establishments (for example, those > the country mean in Fig. 2B) suggests that while biosecurity efforts (all management actions to prevent the introduction and establishment of IAS) may be containing the increase, action must be stepped up to lower the mean annual number of newly established species to reduce downstream invasion impacts. By contrast, flat establishment trends with comparatively low annual numbers of IAS (countries lying to the left of the country-wide mean in Fig. 2B) represent likely greater biosecurity success, i.e. with prevention, early detection and rapid response measures^29^ keeping establishments at consistently low levels.

Accounting for the detection process in the model structure improved establishment rate estimates, particularly with the inclusion in the Sampling model of an independent proxy of observation effort to accommodate non-monotonic changes in effort over time. Consistent with simulated performance differences between models^25^, the Naïve (51% of cases) and Sampling (41% of cases) models best fit the data across most countries (Methods). Comparing the Naïve and best-model slope coefficients across countries reveals the extent to which accommodating the detection process alters rate estimates (Fig. 2C). For countries with positive Naïve model slopes, rate-of-change estimates tended to be larger, reflecting an underestimation of recent establishments attributable to low observation effort in recent years. Conversely, for countries with negative Naïve model slopes, estimates tended to be smaller, reflecting the correction for a lag in IAS detection (Fig. 2C).

The approach taken to counting IAS at the country level is conservative in the sense that it may overestimate local realisation of negative impacts by including species with evidence of negative impact anywhere in their introduced range (Extended Data Table 1) when a negative impact from a particular IAS may not, or may not yet, have been realised in a country in which it is established. Information on local impacts of IAS is especially incomplete^9^, and an alien species becoming invasive anywhere remains a good predictor of its likely risk elsewhere^30^. This approach would nonetheless overestimate the invasion load of a country where this prediction is not realised, biasing upwards the average number of IAS established per year and the rate of change in establishment. Regardless, the country-level inclusion of alien species with evidence of being ‘invasive anywhere’ remains a robust decision for policy reporting purposes, given the precautionary requirement of successful biosecurity^14,15^.

### Ecological explanations for a negative trend

Two ecological explanations for a reduction in the rate of successful new establishments warrant consideration. First, increasing competition from established IAS could theoretically slow the establishment of new ones. However, there is little evidence for IAS outcompeting each other; rather, related alien species and those with similar or facilitatory traits tend to enhance further invasion^31–33^. Second, saturation in the pool of potential new IAS, that is, species whose native ranges intersect with invasion pathways and that have traits making them potentially successful and impactful invaders (estimated here as 16% of established alien species), could explain a declining rate. Although generalist species might be expected to saturate before the broader alien species pool^34,35^, there is no direct evidence for global IAS saturation, and evidence to the contrary for all alien species^1,36^.

### Data completeness and future monitoring

Individual country trend estimates rely on the completeness of national checklists and the accuracy of first record dates (Extended Data Table 1). The expected number of newly established IAS globally for the period ranged from 549 to 651 (Table 1); calculated to show the effect of IAS with no or unknown years of first record on the raw number of establishments between 1970 and 2020 (Extended Data Fig. 3A). In some cases, these years of establishment or first record are unknown, but in many (as shown for the case-study countries in Fig. 2E) the information is available but not yet findable, accessible, interoperable and reusable (FAIR). Expert augmentation did not generally change trend direction for the case-study countries, but it did shift slope coefficients in some cases and substantially increased first record completeness (from 28–89% to 37–94% after augmentation; Extended Data Table 2). This shows that targeted national investment in FAIR data on dates of IAS first record is critical for refining country-level trend estimates and monitoring rates of establishment as an indicator of policy and management success from now on.

Biological invasions are an inherently transboundary phenomenon, necessitating the sharing of information and FAIR data for effective biosecurity across countries^37^. For example, a concerted effort over the last 15 years saw the number of countries with IAS checklists (of any form) rise from 25% to 97% of countries with FAIR checklists for IAS (Extended Data Table 1, top row)^38,39^. A similar effort by countries to improve the completeness of dates of first record (Extended Data Table 1) of IAS will substantially improve country-level estimates of invasion trends for many countries, particularly those not modelled here. Currently, the collation and publication of such data rely largely on the independent efforts of a small number of scientists. Governments and country experts are well-positioned to lead this effort, as shown by the case-study countries where expert knowledge substantially extended the available first records.

### Invasion debt and credit

The concept of invasion debt refers to those alien species introduced to a country but not yet established, widespread, impactful, or managed^40^. Similarly, invasion credit can be considered to be those IAS not yet introduced into a country and where there is the potential to prevent their introduction through biosecurity measures. Although the numbers of IAS globally are in the thousands, most of these species (>80%) remain established in fewer than 10% of countries (Fig. 4A), and of those that emerged between 1970–2020, 80% are established in less than 4% of countries (Fig. 4B). The degree of invasion credit is thus substantial, although stronger for invasive alien animals (93 % in < 10 % countries) than plants (77% in < 10% of countries).

**Fig. 4.**
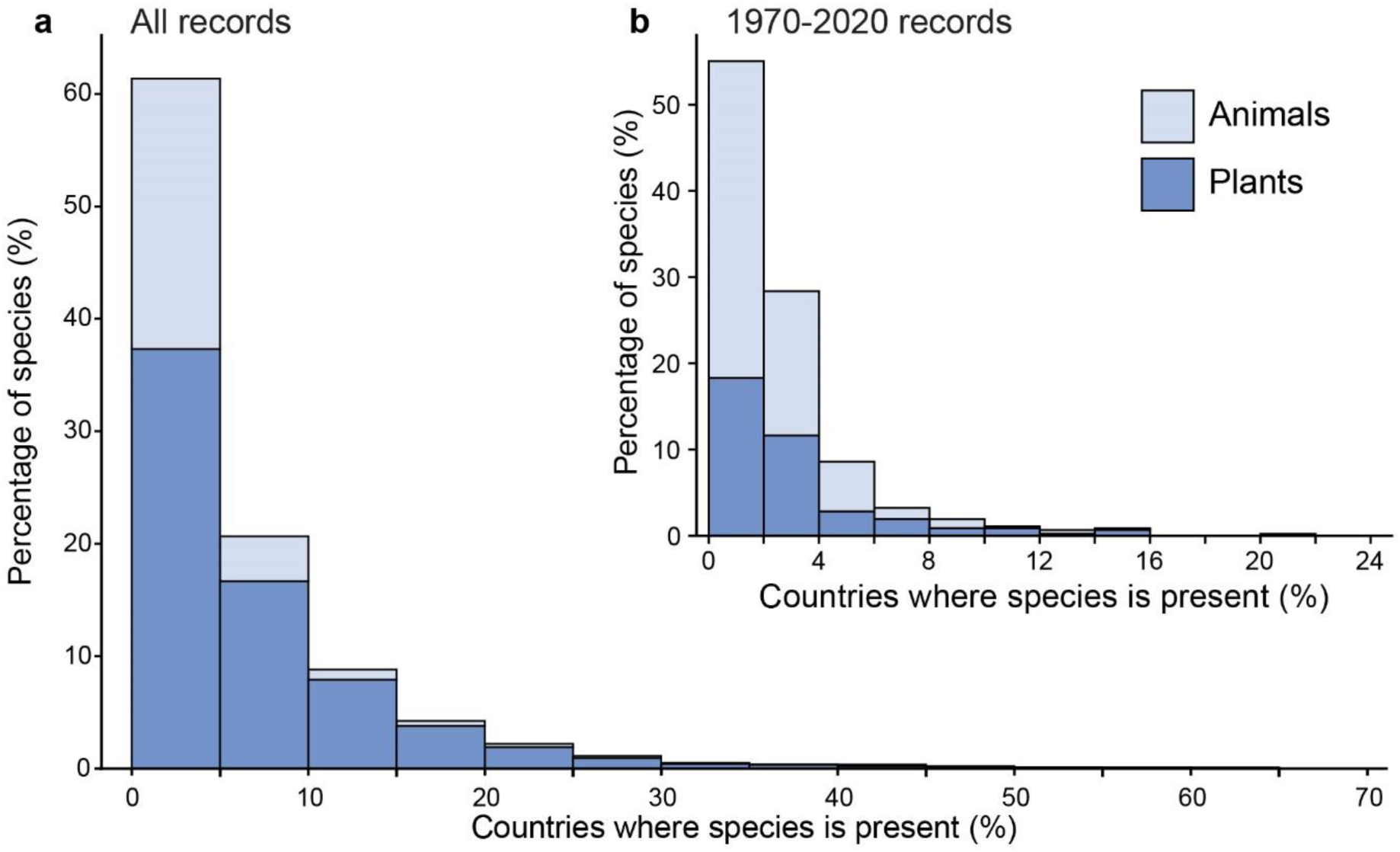
The frequency with which individual invasive alien species (IAS) are found across multiple countries. (**a**) The percentage of countries (and including Antarctica, *n* = 194) occupied by each IAS in the global checklist of established IAS (*n* = 5087), showing that the majority of IAS (>80%) remain established in fewer than 10% of countries. (**b**) The percentage of countries (*n* = 194) occupied by each IAS with earliest first records between 1970 and 2020 (i.e. included in the modelled time-series for the global analysis, *n* = 465), showing that the majority of more recently established species (>80%) are, to date, established in fewer than 4% of countries.

The narrow geographic spread of most IAS (Fig. 4A-B) can thus be considered to be an invasion credit at either global (the number of global IAS not (yet) widely distributed across countries) or country (the number of global IAS not (yet) present in a particular country in which they have the potential to invade) scales. At the global scale, the proportion of IAS that are not yet in a country is a combined function of the success of the country’s biosecurity and a degree of natural protection (including the suitability of its niches to the species invasion, as well as how exposed it is, e.g. how isolated or distant from sources of extant invasions^41–43^). Importantly, invasion credit can be overestimated when there is inadequate investment in surveys and in FAIR data. Here, the global IAS checklist (Extended Data Table 1; Methods) provides a tool for tracking the establishment of known IAS and the emergence of newly recognised high-priority species globally, and for validating their presence or absence in individual countries for biosecurity purposes. The current invasion credit can be preserved through sustained investment in prevention to limit further intercountry spread of the world’s most impactful invaders.

### Target 6 of the Kunming-Montreal Global Biodiversity Framework

Here we show for the first time that the global rate of IAS establishment has slowed over the 50-year period to 2020. Integrating a suite of data and methodological advances at global and country scales (Extended Data Table 1), the evidence directly informs the outcome of Headline Indicator for KM-GBF Target 6, *Rate of invasive alien species establishment*, providing a measurable baseline against which progress toward the target for 2030 can now be assessed. At an estimated 35% reduction in IAS establishments since 1970, progress falls short of the 50% reduction target. We further show a declining trend for one quarter of modelled countries, with an additional 39% with low and stable rates. However, high average annual numbers of IAS in a subset of countries, and substantial scope for established IAS to continue to spread and cause harm within and across countries^2^, means that strategic and ongoing action on IAS remains essential to minimize the rise in IAS impacts on biodiversity and ecosystems.

Although the 35% global decline in the rate of IAS establishments falls short of the 50% reduction called for by KM-GBF Target 6, it nonetheless signifies measurable progress. The result counters the prevailing assumption that establishment rates continue to rise unabated ^6^ and provides a quantitative baseline against which future decadal progress can be assessed. The most plausible explanation for this decline is a positive outcome of the multilateral environmental policy on IAS, which has grown substantially in geographic and invasion pathway coverage since the early 1990s^44^. The management of IAS has evolved from localised and *ad hoc* attempts at control to strategic, comprehensive frameworks targeting the full invasion continuum^45^, in particular pre-introduction pathway management and early detection and rapid response^46^. International cooperation to coordinate pathway management, combined with technological progress that enables more rapid species identification, data capture and knowledge sharing^14^, has also brought about a fundamental shift from reactive to proactive governance of biological invasions over this period.

There are several challenges to estimating invasion trends for policy reporting. For many individual countries progress against Target 6 will be hard to validate; only 43% of countries had adequate data for model estimation (Fig. 2). For the remaining countries, in some cases (∼ 25% of countries) the available data could warrant a qualitative interpretation of the trend^5^, whereas in others the data are too patchy for analysis (∼ 31% of countries). With little consideration of the capacity and readiness of countries to provide robust evidence against this target and its headline indicator, collaborative efforts by researchers, global research networks (e.g., GEO BON) and intergovernmental networks (e.g., GBIF) continue to deliver globally comparable policy-relevant evidence on biological invasions – as an integral component of a global biodiversity observation system^37,47^. With a decade until final reporting on the KM- GBF, progress towards achieving Target 6 is evident at a global scale. A concerted data collection and collation effort by nations over the next decade and beyond would warrant recalculation of the global trend (Fig. 1) across six decades^25^.

### Conclusion

Looking forward, the large invasion credit that persists globally, with the large majority of IAS still established in only a handful of countries, is both a risk and an opportunity. Sustained investment to prevent and record the introduction, establishment and spread of the world’s IAS with greatest impacts on biodiversity, and including the consideration of highly impactful microorganisms^48^, will be critical to minimising future impacts and management costs. The elements needed to enable this, and drawn on to deliver the results here, include FAIR data^23,24^, indicators supported by Essential Biodiversity Variables^47,49^ and the relevant biodiversity data standards to enable interoperability^50–52^. The remaining bottleneck of completeness of first record dates can be addressed by national investment in data mobilisation, using established standards and shared for global benefit through existing infrastructures^51,53^. Closing the knowledge loop between evidence-informed policy and its downstream effects is essential to reduce the harms caused by IAS and to meet the ambition of KM-GBF Target 6.

### Online content

Extended metadata and data are available at https://doi.org/10.5281/zenodo.22022042. Code is available at https://doi.org/10.5281/zenodo.22022073.

## Methods

The data handling and data modelling workflow is represented in Extended Data Fig. 1. The scope of this work encompasses terrestrial and freshwater realms, plants and animals, for the policy-relevant period 1970 to 2020 (technically 51 years inclusive of 1970 and 2020, hereafter “the period”)^54^. Marine invasive alien species were not included because they require a workflow that differs at multiple steps from the one used here. Archaea, fungi, bacteria, chromista, protozoa and viruses were not included. Data were modelled globally (including all countries as well as continental and maritime Antarctica) and separately for all countries that met a data completeness threshold (see Modelling decisions below).

All data handling, modelling, and visualisation was completed using R version 4.4.0^55^ in RStudio version 2023.6.1.524^56^.

## Data handling

Data were obtained and integrated from three primary online sources, with additional data augmentation as described below (Extended Data Fig. 1):

### (1) Invasive alien species (IAS) data

Species flagged as ‘isInvasive’ in the country checklists of the Global Register of Introduced and Invasive Species (GRIIS) (https://cloud.gbif.org/griis)^23^ were used for all countries (most recent version available as of 1 September 2025). GRIIS provides the most comprehensive open and comparable source of information at the country scale for all taxa on the IAS (those with negative impacts, definition in Extended Data Table 1) established in a country, and several countries use and update their GRIIS checklists as a national data resource^23,39^. As of September 2025, 191 countries had findable, accessible, interoperable and reusable (FAIR) GRIIS checklists available through the GBIF Integrated Publishing Toolkit (IPT Version 3.3.2)^57^ that could be accessed and processed through a bespoke automated workflow (see Harmonisation and integration below).

Two countries do not have FAIR GRIIS checklists available through the GBIF IPT, as these host bespoke IAS checklists through different platforms and contain different data, meaning they could not be harmonised and integrated using the automated workflow. These two countries, and an IAS checklist for continental and maritime Antarctica^58^, were not considered individually, but the bespoke checklists were manually integrated with the output of the automated workflow to produce the global IAS checklist (see Data augmentation below). Three additional countries had available checklists hosted by the GBIF IPT, but formatting of these lists was not interoperable with the automated workflow due to missing data columns and these countries were not included.

### (2) First records data

The year of establishment of each species for each modelled country (definition in Extended Data Table 1) was taken as the year of first record from the SInAS dataset of alien species occurrences, which includes first record dates (version 3.0, accessed 15 August 2025)^59^.

The year of first record of an IAS is the estimate of the year of establishment of a species (Extended Data Table 1). This estimate is understood to be affected by the lag in the detection process (the time between actual establishment and first detection, data capture and reporting) and the increasing likelihood of detection as the invasion proceeds^19,36^ (Extended Data Table 1). The rationale for our modelling approach (see Trend estimation models below) is to accommodate these effects in rate estimation^25^.

### (3) Observation effort proxy

The third primary open data source was the number of new biodiversity occurrence records per year in GBIF. Total annual counts of animal and plant occurrences added to GBIF between 1970 and 2020 (inclusive) were collated for each country using Structured Query Language (SQL) searches through the *occ_download_sql* function of the *rgbif* package (version 3.8.3)^60^. For each country, only those occurrences where both the top-level division from the GADM database (*level0gid* SQL column) and the 2-letter country code (as per ISO-3166-1; *countrycode* SQL column) matched the country details were included, which served to exclude records labelled with a specific country name but occurring outside the country’s land borders (i.e., marine records). Occurrences where kingdom was labelled “*plantae*” or “*animalia*” were included when the basis of record label was any of: “*observation*”, “*living specimen*”, “*material sample*”, “*human observation*”, “*machine observation*”, or “*occurrence*”. Additionally, occurrences were filtered to include only those where the *occurrenceStatus* label was “*present*”, and there was no discernible coordinate issues associated with the record. Observation effort proxy data was downloaded for all countries between 15-17 October 2025, and for the global augmented models, separately for plants and animals, on 5 March 2026.

These data were used in the Sampling model (see Trend estimation models below) as a proxy of IAS observation effort. Very few countries have data and/or open data on observation effort specifically aimed at detecting IAS^45^, meaning that broadly comparable taxon- or region-specific effort estimates specifically focused on IAS are impossible to derive. Therefore, we use the count of new biodiversity records per year in GBIF as a simple, meaningful, comparable and globally generalizable proxy for IAS observation effort. Use of this proxy assumes that the differences in annual number of new biodiversity records over time reflect differences in observation and reporting activity, including IAS, and that an increase in this number reflects an increase in the likelihood of recording new IAS.

We modelled the temporal change in this effort proxy between 1970 and 2020 with generalised linear models, using year as the predictor variable for each country and globally (with a negative binomial distribution, unless significant over- or under-dispersion was detected, in which case a quasi-Poisson distribution was used). These models demonstrated significant growth in biodiversity records since 1970 globally, with a positive slope and P<0.001 (for all records, animals and plants) (see accompanying data). This provides general support for model assumptions that the likelihood of IAS detection has similarly increased with time.

There is no global consensus on how to best estimate spatiotemporal variation in observation effort. Metrics include the count of records for a particular region or taxon, where possible accounting for actual expected presence^61,62^. At the resolution of countries globally and across a wide range of taxa, we suggest that the total count of new biodiversity occurrence records provides an intuitive and highly scalable proxy for the overall observer activity as relevant for IAS. All else being equal, we expect countries with fewer overall records reported to have an accordingly smaller chance of reporting IAS, and vice versa^5,13^. Alternatives could include using GDP, as this may reflect changes in research capacity and funding over time^13,63^, or changes in the numbers of scientific publications in relevant fields^22,64^. However, these metrics are either less directly related to observation effort (GDP) than new biodiversity records, and/or using economic and trade-related variables confounds information on drivers of invasion with those accounting for the effect of observation effort on first records^13^ (Extended Data Fig. 3).

### Harmonisation and integration

#### Country scale analysis

The SInAS workflow (version 2.0.1)^65^ was used as the first step to standardise and harmonise taxonomic information across each individual country’s IAS checklists (primary online source 1 above) and associated first record data (primary online source 2 above). The SInAS workflow facilitates the standardisation of terminology across datasets, including the harmonisation of country nomenclature, taxonomic information, and event dates. Following the initial application of this workflow, several country and species name records remained unresolved. To address this, the workflow was extended with an additional quality control stage, incorporating both automated and manual harmonisation procedures to resolve discordant location and taxonomic entries.

Unresolved geographic records were addressed through a manual correction step, in which ambiguous or unmatched location strings were reviewed and assigned to the appropriate country.

Taxonomic names that could not be harmonised automatically were subjected to a structured manual correction process. Each unresolved checklist name was queried against the GBIF taxonomic backbone using the *name_backbone_verbose* function from the *rgbif* package^60^. The resulting match outcomes were evaluated according to a decision framework, with the following rules applied:

- Hybrid taxa, identified by the presence of “x” (upper or lowercase) in the original name string, were retained verbatim without modification.
- Where the GBIF matching algorithm returned a fuzzy match, the associated taxonomic metadata were reviewed and corrected manually.
- For names returning an exact match or a match at a higher rank – provided the matched rank was at species or genus level – the resolved name was accepted as valid. Names matched only at higher taxonomic ranks were subject to further manual correction.
- Names flagged by GBIF as doubtful were manually corrected on a case-by-case basis.
- Where a matched name was returned as a synonym or homotypic synonym with an exact or higher-rank match, the synonym was accepted and adopted accordingly.

Resolved names were reintegrated into the final analytic dataset.

#### Global scale analysis

Once harmonised and integrated, IAS and first records data were available for each country. and These country lists were further integrated to generate a global dataset of IAS, with associated first record dates for each country where these IAS are present (where available). The global checklist was then further augmented (see Data augmentation below).

## Data augmentation

### Country scale analysis

Five case study countries were used to ground-truth the workflow approach: Australia, Benin, Nepal, Norway and Poland. Country experts (see acknowledgements, including authors of this manuscript) were consulted on the data and model interpretation in each case. Experts were asked to add missing first records where available, edit existing first records where more accurate data was available, and to verify or edit the inclusions of established IAS. For each of these countries there were therefore two datasets, one derived from the automated workflow and the augmented dataset reviewed by the expert (Extended Data Table 2).

### Global scale analysis

The data collated for each of the 191 countries from the automated workflow were augmented using additional available datasets of IAS first records at the global scale and for selected countries and regions. Specifically, while IAS checklists for the United States, Belgium and continental Antarctica could not be processed using the automated workflow, openly accessible lists of IAS for these regions were manually integrated with the outputs from the automated workflow to augment the global checklist^58,66,67^. Additionally, for the five case study countries, changes to the IAS and introduction dates in the expert-derived augmented lists were integrated into the global data.

Finally, IAS introduction dates were compared between the global checklist and the Global Invasive and Alien Traits And Records (GIATAR) dataset^68,69^. First records for country- species pairs (specific species in specific countries) were compared across datasets, and where earlier introduction dates were identified in the GIATAR dataset (*n* = 1826 unique species), that introduction date was adopted for the global dataset, to consolidate additional potential sources of first records included in the GIATAR dataset that had not previously been captured by the automated workflow. A total of 5087 unique invasive species were included in the final global IAS checklist.

### Model-ready and formatted time-series data

As a result of the data handling workflow, three distinct sets of harmonised and integrated IAS datasets were utilised for further analysis:

i. Country-scale IAS checklists for 191 countries, generated through the automated workflow;
ii. Augmented country-scale IAS checklists for the five case-study countries, based on updating the original checklists through consultation with country experts (the “augmented” country checklists);
iii. A global-scale IAS checklist, based on the collated country-scale checklists with manual adjustments to or addition of country IAS lists (including Antarctica), and integration of further sources of first record years.

These checklists were used to create time-series of IAS introductions between 1970–2020 for global-scale analyses, overall and split by plants and animals, as well as for country-scale analyses, for those countries considered to have adequate data for modelling (see Modelling decisions below).

For individual countries, IAS with establishment dates within the policy-relevant period were summed per year across the 50-year time series, and these data were integrated with the observation effort proxy annual values (primary online source 3 above) across the same period. For the case-study countries, datasets were created for both the original (integrated open) and augmented data, and both were included in further analysis to allow comparison of results (Extended Data Table 3).

At the global scale, only the earliest establishment date recorded for each IAS in any country was considered. The global time series data therefore reflects, for each year, the number of IAS that first became or were recorded to be established in any country (outside their native range) between 1970–2020.

For the global time series data, to ensure the observation effort proxy was taxon-specific when modelling plants and animals separately, new SQL searches were performed for GBIF occurrence records for plants and for animals from all included countries (and Antarctica) and these plant and animal occurrence values were summed per year for the overall global-scale analysis.

All model-ready time series datasets were then assessed prior to analysis to consider the number and distribution of IAS establishments over the 50-year period, with specific criteria applied to determine modelling approach (see Modelling decisions below).

## Data modelling

### Trend estimation models

Four models were used to estimate the trend direction (sign of the slope coefficient, β_1_), rate of change across years (value of β_1_) and rate of IAS establishment (average number of IAS established per year): a Naïve model and a family of three related models, i.e. the Solow and Costello, Constant Detection, and Sampling models, as outlined with statistical and ecological rationales in Buba et al. ^25^. Modelling of the Solow and Costello family of models was conducted using the *snc* function of the *alien* package in R^70^.

These four models make different ecological assumptions about the detection of new IAS, and based on simulated evaluation of each model, each performs better under a different set of data and model parameter estimate conditions (for full rationale and formulation see ^25^).

#### (1) Naïve model

This model assumes that there is perfect detection of IAS, i.e. there is no temporal lag between the year of establishment of a new IAS, and the year (*t*) of its detection and first record.

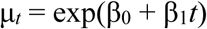

Where β_0_ defines the initial introduction rate and β_1_ defines the change in the introduction rate with time. The AIC values of this model were compared (ΔAIC) with a constant rate null model μ*_t_* = exp(β₀). The Naïve model tends to perform better (i.e. has lower estimated bias) than the other three models when the mean baseline annual detection is high (i.e. more closely meeting the assumption of perfect detection), including in short time-series and when detection probability varies non-monotonically^25^.

#### (2) Solow and Costello model

This model was developed originally to parametrise and estimate the variability in alien species establishment time series caused by the lag in the detection process^19^, i.e. here the lag between species establishment, detection and reporting or data capture. In this model, the number of species first recorded in year *t* follows a Poisson distribution. The model assumes a monotonic increase (with saturation) in the probability of detection of IAS over time, driven by (i) observation effort and (ii) exponential growth in species abundance making them more likely to be detected (see ^25^ for full model formulation and rationale). The model maximum likelihood (ML) function estimates five parameters related to the introduction and discovery process: β_0_ and β_1_, parametrising the introduction process as in the Naïve model, and three parameters related to the discovery process: γ_0_, γ_1_, and γ_2_, which describe the baseline observation effort, the increase in observation effort with time, and the increase in detectability of newly established IAS with increasing post-introduction abundance (assumed to follow an exponential distribution), respectively. This model is often best supported in cases where both the mean introduction rate is high, and the baseline detection probability is high and varies monotonically^25^.

#### (3) Constant Detection model

This model is a modified form of the Solow and Costello model with a reduced number of parameters. The ML function estimates three of the same parameters as the Solow and Costello model (β_0_, β_1_, γ_0_), but to avoid potential overparameterization for shorter or more data-sparse time-series, assumes that detection of IAS is imperfect but constant across the period (and therefore does not include estimates of γ_1_ and γ_2_). This model is most commonly the best supported model when baseline annual detection (γ_0_) is low and changes monotonically, but can perform poorly (i.e. has high estimated bias) when detection is non-monotonic^25^.

#### (4) Sampling model

This model was developed to address the temporal and spatial variation in the likelihood of detecting IAS and incorporates external data on IAS observation effort. It assumes there is a non-monotonic, variable observation effort for new IAS establishments that is parametrised using an observation effort proxy (i.e., the number of new biodiversity records per year, primary online source 3 above). Detection probability is therefore not assumed to vary linearly with time and is instead determined by the deviation from mean observation effort in each year. Estimated model parameters include β_0_ and β_1,_ as in each of the previous models, as well as ζ_0_ and ζ_1_, which correspond to the baseline annual detection in the absence of changing observation effort, and the change in detection with change in the observation effort proxy, respectively. This model is most often best supported where the detection trend is non-monotonic and variable over the time-series, and the baseline annual detection (ζ_0_) is low to medium^25^.

## Modelling decisions

Fifty years is considered the minimum length suitable for estimating model parameters with an acceptable degree of and variability in bias in the Solow and Costello family of models^25^. The results were shown to be robust to the 1970–2020 time series bounding.

First, the time series records extend beyond the selected period, indicating that there is no strong boundary effect. Second, the trend outcome is largely consistent with different series’ lengths and decadal inclusion (when modelled using the Naïve model, due to the shorter time-series lengths; Extended Data Fig. 2).

For the country-scale analysis, summary statistics of each country’s harmonised IAS checklist were used to determine the modelling approach taken for each country. Our intention was not to directly compare the results of individuals countries but rather to examine variation across them based on best model selection in each case (see Model selection below). Two measures to determine the paucity and/or sparsity of IAS data were used:

i. The proportion of IAS listed in each country’s GRIIS list that were successfully harmonised to the SInAS database and where a year of establishment/first record was identified (hereafter the “first record completeness”), and
ii. The number of years within the policy-relevant period where at least one IAS

establishment was recorded (hereafter the number of “non-zero years”).

First record completeness was assumed to align with the availability and reliability of IAS establishment date data within a country, as the number of species that may have been introduced within the policy-relevant period for countries with very low first records completeness cannot be readily determined. The number of non-zero years across the time series, while not related to data completeness, nonetheless constrains model options when low. The Solow and Costello family of models requires ML estimation of 3-5 parameters, and countries with few or unevenly distributed/clustered records of IAS establishments (i.e., few non-zero years) within the policy-relevant period were unable to have models reliably fitted to their time-series data due to potential over-parametrisation.

Countries were categorised into different modelling groups based on criteria related to both first record completeness and non-zero years. All countries with fewer than 15 non-zero years across the policy-relevant period (i.e., fewer than 15 years within which at least one IAS establishment was recorded) were considered too data sparse to model and were excluded from further analysis. Similarly, countries with < 25% first records completeness (where the year of establishment of more than 75% of IAS present in a country’s GRIIS list was unknown) were not considered for further analysis. For countries with between 15 and 25 non-zero years, and greater than 25% first record completeness, the best working model was selected from the Naïve model, Constant Detection model and Sampling model (see Model selection below), but the Solow and Costello model, which estimates five parameters, was not included. Finally, for countries with more than 25 non-zero years (i.e., at least one IAS establishment was recorded in half of the years across the policy-relevant period) and with greater than 25% first record completeness, model selection to identify the best working model included all four model options (see Model selection below).

The global time-series data met the modelling decision criteria for all four models to be run and compared when selecting the best working model. For the country-scale analysis, nearly half of countries met the inclusion threshold criteria to be modelled (43%, *n* = 85). Of these, model selection from all four potential model options was possible for 43 countries, and for the remaining 42, model selection was based on comparing the Naïve, Constant Detection and Sampling models only. Note that for the country analysis the decision to model a country is based on adequate data (a sufficient percentage of IAS with first record and number of non-zero years) to run and interpret the model. Nonetheless, when interpreting the policy-relevance of the model outcome for each country (trend and mean annual number of IAS), the percentage completeness of first records must be further taken into account.

## Model selection

Model selection criteria were informed by the simulated model evaluation results presented in Buba et al. ^25^ that identify the model parameter conditions under which each model is likely to perform best, and predicated on the parameter value interpretations outlined in Buba^70^. The AIC of the Naïve model is not comparable with the AIC of the Solow and Costello family of models as they have different likelihood functions, and as such, the ΔAIC of the Naïve model was calculated from a null (intercept-only) model. AIC was compared across the Solow and Costello family of models and considered as part of the model selection process.

Once selected, the best working model was used to assign each country a trend category (negative, flat, positive trend in IAS establishment), and to determine the slope estimate. Non-significant Naïve models were considered flat, and a flat trend was assigned to other models using the standard error of the slope coefficient, and the cumulative distribution function of the probability of it being different from zero. Agreement across multiple models increases confidence in the result beyond that of any single model, providing additional support^71^.

## Observed versus expected numbers of IAS

The ‘observed’ number of IAS is the number of IAS at the global or country scale derived from the workflow, including the augmented data where relevant. The ‘expected’ number of IAS for the period 1970–2020 was calculated because a first record date was not available for all IAS, which meant that a species could not be assigned as introduced during the policy-relevant time period or not. This expected number was derived assuming a random temporal distribution for all of the IAS with missing first records across the observation period (before and after 1970), and then subsequently adding the randomly allocated first record dates that fell between 1970–2020 to the observed number, and using Wilson score test to calculate 95% confidence intervals. The relationship between the observed and expected number of IAS is shown in Extended Data Fig. 3A. For the five case study countries, the expected number of IAS was calculated from the observed number derived from the automated workflow, rather than from the augmented list, and was then compared to the observed number of IAS in the augmented data. The augmented data from case study countries fell within the estimation range for some countries, suggesting that in these cases the augmented data provide a reasonable approximation of numbers of IAS (Extended Data Table 2). Note that these expected numbers of IAS are not comparable with the model-derived estimates.

## Data exploration

A range of relationships between data properties and model estimates were examined for further insight on geographic and other patterns in the data (Extended Data Fig. 3B-F). To assess whether country size or per-capita GDP may influence the change in rate of IAS establishment, measures of total country land area (km^2^) and per-capita GDP in 2020 (in current US$) were sourced for modelled countries (*n* = 85)^72,73^. Plots depicting the relationships between country properties and country-level IAS statistics (observed number of IAS, mean IAS, and slope coefficient (β_1_) of the best-selected model trend) are shown in Extended Data Table 3.

## Data availability

The datasets generated and analysed during the current study are available at Zenodo (https://doi.org/10.5281/zenodo.22022042).

## Code availability

The code used for data analysis is available at Zenodo (https://doi.org/10.5281/zenodo.22022073)

## Acknowledgments

We thank Ezekial Buba, Shyama Pagad and Hanno Seebens for their foundational work enabling this research. Hanno Seebens kindly commented on a draft version of the manuscript. Thanks to Noel Cressie for discussion on the research approach, Ane Marlene Myhre for reviewing the Norwegian data, and Andrew Rodriguez for assistance with GBIF biodiversity record data.

## Author contributions

Conceptualization: MAM, JB, MW, WJ, SA, RTM; Data curation: RTM, SA, JCG, BBS, WS, MAM; Formal analysis: RTM, SA, MAM; Investigation: MAM, RTM, SA; Methodology: MAM, RTM, SA, WJ, RIL, BRS, JB; Visualization: RTM, MAM, BRS; Funding acquisition: MAM; Writing – original draft: MAM, RTM; Writing – review & editing: MAM, RTM, SA, JB, JCG, WJ, RIL, BRS, BBS, WS, MW.

## Funding

This work was supported by Monash University and ARC SRIEAS Grant SR200100005 Securing Antarctica’s Environmental Future (M.A.M., R.M., S.A., R.I.L.), as well as by the ARDC Nectar Research Cloud, a collaborative Australian research platform supported by the Australian Research Data Commons (ARDC). The Group on Earth Observations Biodiversity Observation Network (GEO BON) and the Global Biodiversity Information Facility (GBIF) provided in-kind support.

## Competing interests

The authors declare no competing interests.

---

**Extended Data Table 1.** Concepts and operational definitions of model variables and data outputs at global and country scales - used in estimating the establishment of invasive alien species (IAS). (**a**) ^6^, (**b**) ^23^, (**c**) ^36^.

| Operational definitions | Global analysis | Country analysis |
| --- | --- | --- |
| **Invasive alien species** |  |  |
| The subset of established alien species that spread and have a negative impact on biodiversity, local ecosystems and species.<sup>a</sup> | Any alien species established and considered to be an IAS anywhere.<sup>b</sup><br>**Data:** Global IAS Checklist. | Any alien species established in that country and considered to be an IAS in that country of elsewhere.<sup>b</sup><br>**Data:** Country IAS checklists. |
| **First record** |  |  |
| The year of first record <sup>c</sup> of each IAS is taken as the year of first establishment of an IAS, with the knowledge that this year is likely later than the true year as a result of the detection process (the lag time between when a species establishes and its first record). | The year of first record of a species across all countries (including Antarctica) that falls within the policy-relevant period 1970-2020.<br>**Data:** Global first records (1970-2020). | The year of the first record of a species in the country, where this year falls within the policy-relevant period 1970-2020.<br>**Data:** Country first records (1970-2020). |
| **Detection process** |  |  |
| The detection process includes the observation, recording, verification, data capture and publication of the IAS record.<br>This process results in a lag between the date of alien species establishment and the date of its first record, with the extent of the lag influenced by both observation effort and the likelihood of detection that increases after establishment as the IAS population grows and spreads<sup>19</sup>. |  |  |
| **Trend and rate of establishment** |  |  |
| The trend (positive, flat or negative) and rate of change in establishments of new IAS for the policy relevant period 1970-2020. | The trend and rate of IAS establishment globally across the policy-relevant period 1970-2020.<br>**Data:** Global IAS checklist and first records. | The trend and rate of IAS establishment in each country across the policy-relevant period 1970-2020.<br>**Data:** Country IAS checklist and first records. |

**Extended Data Table 2.** Comparison of the original, integrated open data and the outcome after augmentation by country experts for the five case study countries, demonstrating the scope for increased mobilisation and curation of invasive alien species (IAS) data at the country level.

| Statistic (1970–2020) |  | Australia | Benin | Nepal | Norway | Poland |
| --- | --- | --- | --- | --- | --- | --- |
| **Number of IAS, observed (expected)** |  |  |  |  |  |  |
| Original data number |  | 166 | 17 | 30 | 95 | 96 |
| Expected number |  | 187 | 60 | 106 | 186 | 172 |
| [95% CI] |  | [162,216] | [48,63] | [81,1310] | [155,221] | [144,204] |
| Augmented data number |  | 196 | 26 | 58 | 146 | 89 |
| **First record completeness** |  |  |  |  |  |  |
| Original data |  | 89% | 28% | 28% | 51% | 56% |
| Augmented data |  | 94% | 56% | 37% | 91% | 93% |
| **Nature of Augmentation, between 1970–2020 (total)** |  |  |  |  |  |  |
| Species | Added |  |  | 30 (31) | 2 (4) | 22 (33) |
|  | Removed (considered not established) |  |  |  | 45 (428) | 29 (235) |
| First records | Added | 30 (78) | 12 (18) | 16 (16) | 95 (262) | 23 (161) |
|  | Revised to be outside of 1970–2020 |  | 3 | 18 | 2 | 23 |
|  | Revised within 1970–2020 |  |  | 3 | 24 | 10 |

**Extended Data Table 3.** Comparison of the model results from models produced using the time-series of original, integrated open data and data augmented by country experts for the five case study countries, demonstrating the effect of increased mobilisation and curation of invasive alien species (IAS) data by countries on establishment trend and rate estimates. Best-selected models for each country and data source are in bold.

| **Data** | **Australia** | **Benin** | **Nepal** | **Norway** | **Poland** |
| --- | --- | --- | --- | --- | --- |
| **Naïve model, $\beta_1$ (s.e.)** |  |  |  |  |  |
| Original data | -0.037 (0.006) | **0.018 (0.017)** | **0.027 (0.013)** | **-0.0002 (0.007)** | -0.005 (0.007) |
| Augmented data | -0.037 (0.005) | **-0.018 (0.014)** | 0.001 (0.009) | 0.017 (0.006) | 0.025 (0.008) |
| **Sampling model, $\beta_1$ (s.e.)** |  |  |  |  |  |
| Original data | **-0.023 (0.007)** | -0.080 (0.130) | -1.945 (4.101) | 0.0021<br>(0.0010) | **-0.067 (0.031)** |
| Augmented data | **-0.025 (0.006)** | -0.018 (0.033) | **0.002 (0.010)** | **-0.007 (0.008)** | **0.027 (0.008)** |
| **Trend** |  |  |  |  |  |
| Original data | Negative | Flat | Positive | Flat | Negative |
| Augmented data | Negative | Flat | Flat | Flat | Positive |
| **Mean IAS establishments per year (best model) (s.e.)** |  |  |  |  |  |
| Original data | 3.255 (0.240) | 0.333 (0.012) | 0.588 (0.033) | 1.863 (0.006) | 1.883 (0.104) |
| Augmented data | 3.843 (0.284) | 0.510 (0.019) | 1.137 (0.009) | 2.863 (0.192) | 1.745 (0.093) |

**Extended Data Fig. 1.**
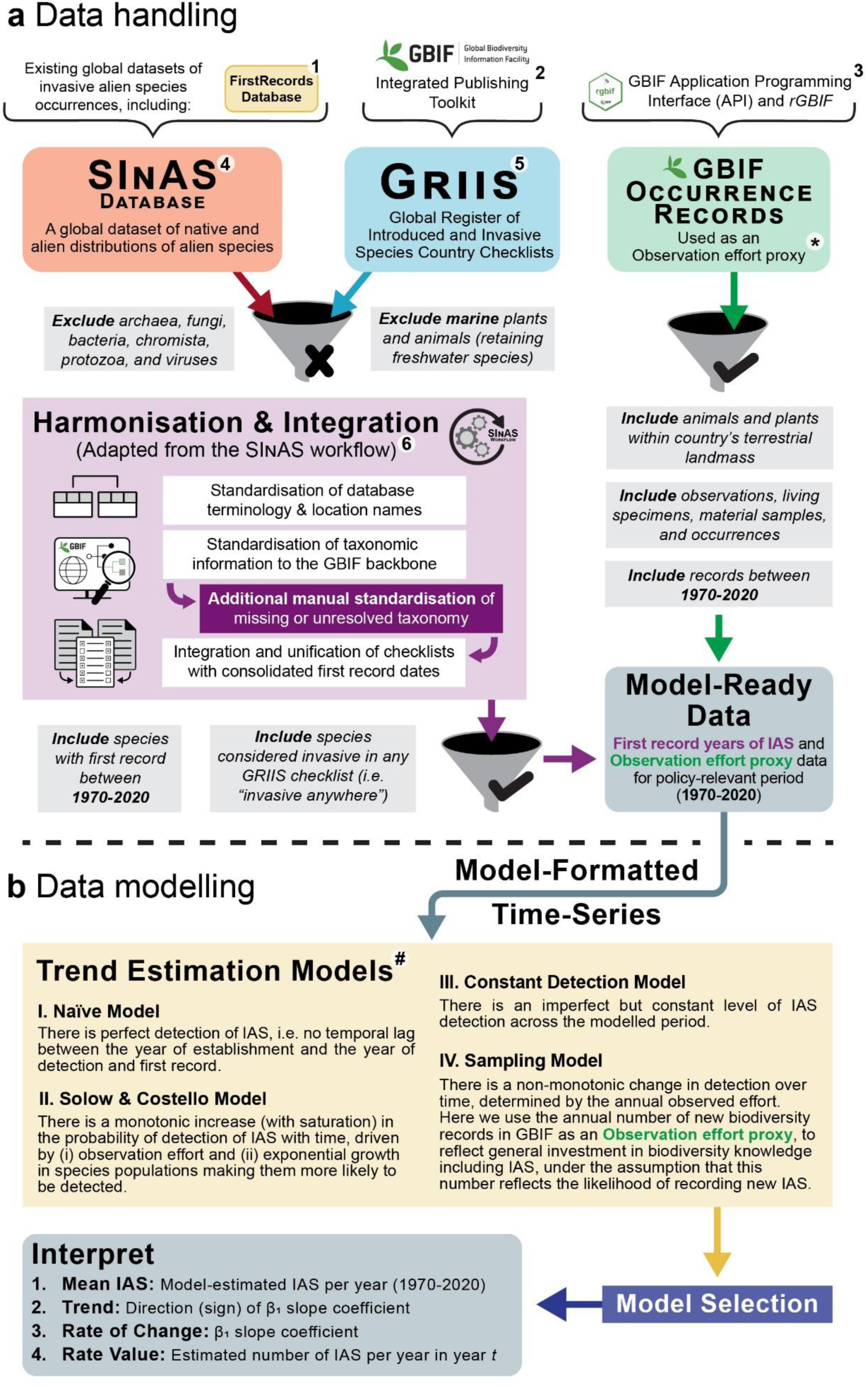
Workflow established to estimate the rate of invasive alien species (IAS) establishment at global and country scales. The (**a**) data handling and (**b**) data modelling steps integrate a suite of methodological advances in biodiversity informatics^51^, findable, accessible, interoperable and reusable (FAIR) data^23,24^, and trend estimation^25^. The workflow has automated and manual components, and italicised text in grey boxes represents data management decisions applied during workflow. \***Observation effort proxy** - The annual number of new biodiversity records used to reflect general investment in biodiversity knowledge including IAS; an increase in this number is assumed to reflect an increase in the likelihood of recording new IAS (used in the Sampling model only). ^#^**Trend Estimation Models -** overview of the four exponential models used and their respective assumptions about the detection process (Methods). 1) ^74^ 2) ^57^ 3) ^60^ 4) ^59^ 5) ^75^ (Methods) 6) ^65^. For further details on each of the workflow steps, see Methods.

**Extended Data Fig. 2.**
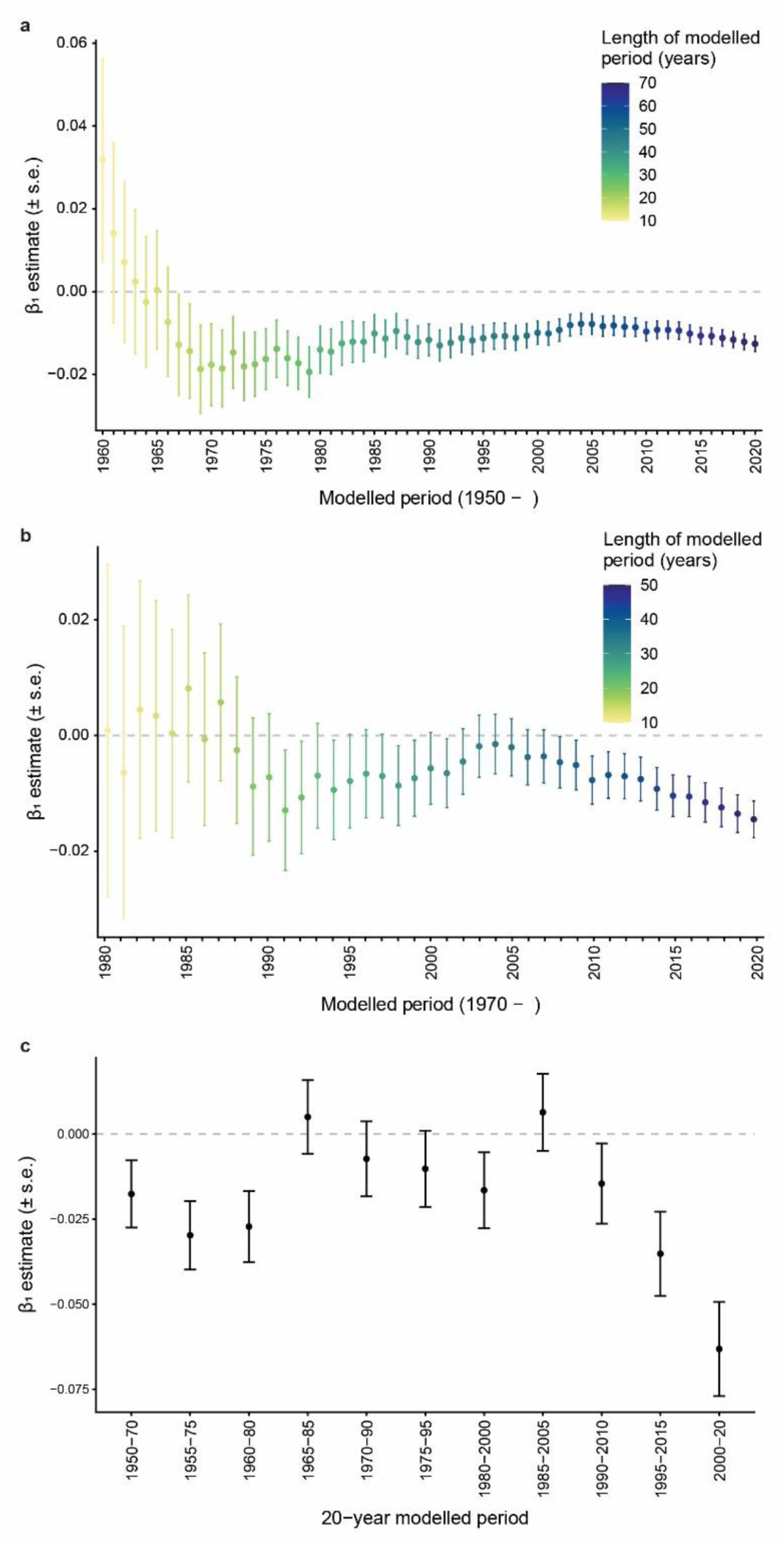
To assess the potential effect of the timing and bounding of the selected period for analysis (1970–2020) on the establishment trend, we examined the change in Naïve model slope coefficients (β_1_, s.e.) of (**a**) successive time-series’, starting two decades earlier, initially for a modelled period of ten years (1950–1960), and then increasing the length of the modelled period by one year until the final modelled period of 1950–2020, (**b**) successive time-series’, starting within the policy-relevant period at 1970, initially for a modelled period of ten years (1970–1980), and then increasing the length by one year until the final modelled period of 1970–2020, and (**c**) moving windows of 20-year time-series, starting with 1950–1970 and shifting by 5 years for each modelled period. These results support the validity of the negative slope coefficient for the global trend, regardless of temporal bounds.

**Extended Data Fig. 3.**
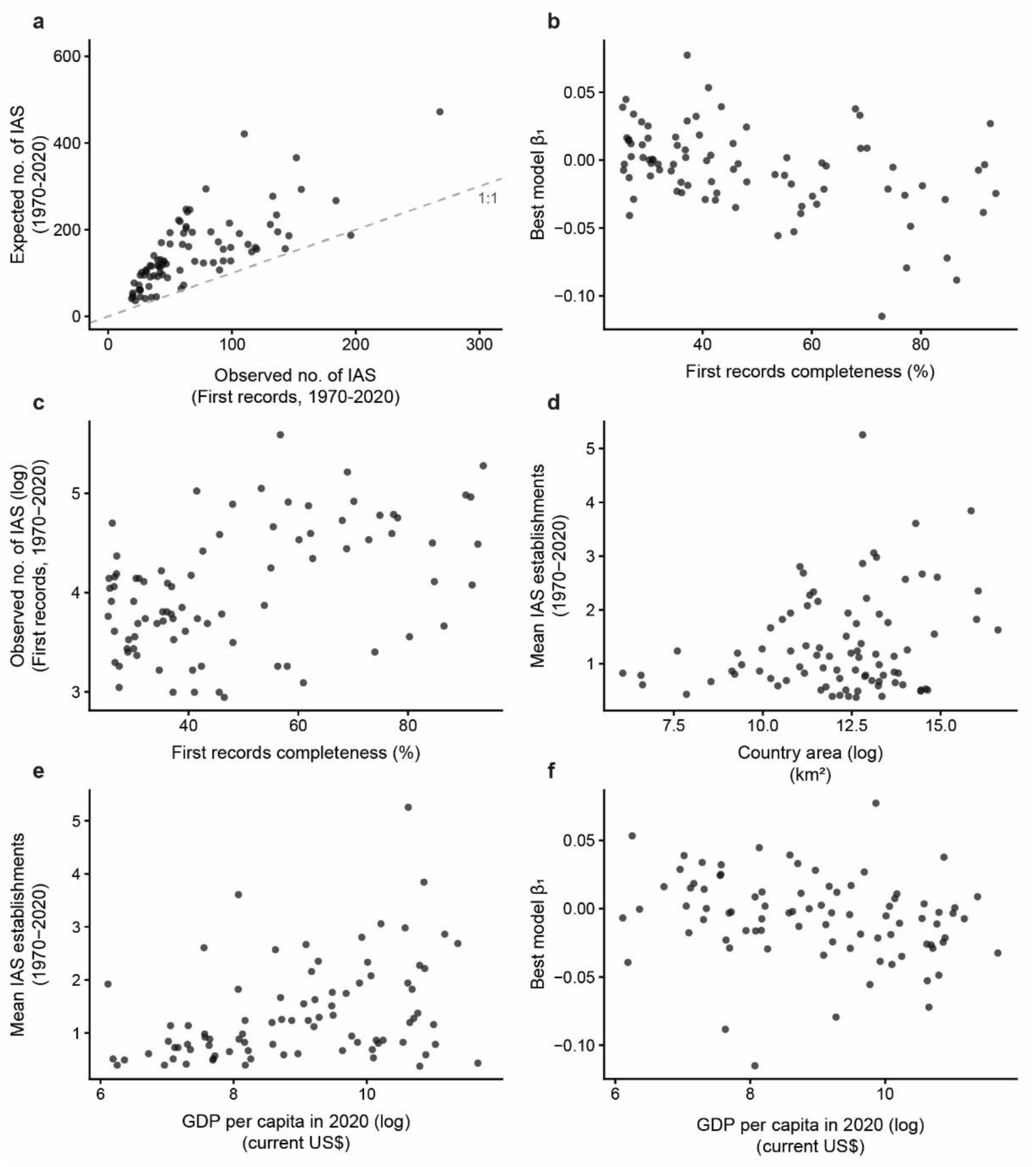
Relationships between country-level IAS statistics (observed number of IAS, mean IAS and slope coefficient (β_1_) of the best-selected model trend) and country properties for the period 1970-2020 including (**a**) expected number of new IAS over the period per country (expected number calculated assuming a random temporal distribution of IAS without first record dates before and after 1970, see text); (**b, c**) first records completeness for the country; (**d**) country area, and (**e, f**) per capita Gross Domestic Product (GDP) in 2020 (log transformed; current US$). Country area data was sourced from ^72^ and GDP data for 2020 was sourced from ^73^. The data include the modelled countries (*n* = 85; in E and F, *n* = 84).

## Notes

### Competing Interest Statement

The authors have declared no competing interest.

